# Structure of the histone acetyltransferase NuA4 complex

**DOI:** 10.1101/2022.07.11.499577

**Authors:** Liting Ji, Lixia Zhao, Ke Xu, Huihan Gao, Yang Zhou, Roger D. Kornberg, Heqiao Zhang

## Abstract

NuA4, one of two major histone acetyltransferase complexes in *Saccharomyces cerevisiae*, specifically acetylates histone H4 and H2A, and therefore loosen the histone-DNA contacts, resulting in increased transcriptional activity. Here we present a near-atomic resolution structure of NuA4 complex through cryonic electron microscopy. The structure comprises six subunits and a total of 5,000 amino acids, within which Eaf1 and Eaf2 twist against each other and span from Actin-Arp4 module to the platform subunit Tra1, forming the backbone of NuA4. The locations of four missing components including Yaf9, and Esa1, Yng2 and Eaf6 of Piccolo module were indicated based on the structure. Intriguingly, our biochemical data showed that NuA4 doesn’t bind nucleosome *in vitro*, whereas possesses strong histone acetyltransferase activity. Its Piccolo module, which has long been regarded as the catalytic module of NuA4, acetylates the nucleosome substrate at almost the same efficiency, but is capable of binding nucleosome. Combining the structural and biochemical data give rise to a model in which the entire NuA4 adopts an auto-inhibited conformation, and undergoes structural rearrangements upon substrate and co-factor additions, restoring its capacity of nucleosome-binding and activity of hyper-acetylation.

## Introduction

In eukaryotes, the genetic information is stored in chromatin, instead of naked DNA (1). The nucleosome, a basic unit of chromatin formed by histone octamer and approximately 147-bp DNA wrapping around it (2), serves as transcription barrier for almost all protein-coding genes (3). Post-translational modifications (PTMs), which mainly occur on the histone tails, influences the chromatin context and therefore regulates transcription (4). Histone acetylation is one of the well-studied PTMs that plays important roles in gene regulation and maintenance of genome integrity (5). Acetylation of histone tails neutralizes the positive charge of lysines in histones, weakening the interaction between histone proteins and negatively charged DNA, and unwinding compacted DNA from histone octamer to promote transcription (5). Histone acetylation level is balanced by histone acetyltransferases (HATs) and histone deacetylases (HDACs) (6). NuA4 (Nucleosome acetyltransferase of H4) and SAGA (Spt-Ada-Gcn5-Acetyltransferases) complexes are two major HATs in *S. cerevisiae* (7). Previous studies demonstrate that NuA4 preferentially acetylates the histone proteins H4 and H2A. In addition to histone tails, NuA4 also acetylates other approximately 250 non-histone substrates, serving as transcriptional coactivators to regulate a variety of cellular processes, including DNA repair, cell cycle progression, and chromosome stability, etc. (8-10).

NuA4, a 13-subunit complex with a molecular weight of 1.04 MDa, shares the largest subunit Tra1 with SAGA, whose major substrate is histone H3 (11). NuA4 also shares four subunits Eaf2, Arp4, Actin and Yaf9 with SWR1, corresponding to DAMP1, BAF53a, Actin and YEATS in TIP60 (10). Therefore, the mammalian TIP60 complex has been regarded as a fusion form of NuA4 and SWR1 in *S. cerevisiae*. The Piccolo module, comprising Epl1, Eaf6, Yng2, and the catalytic subunit Esa1, is responsible for direct acetylation of nucleosome (12). The TINNTIN module, consisted of Eaf3, Eaf5 and Eaf7, recruits the phosphorylated RNA polymerase II, and therefore promotes its progression (13); Previous biochemical studies has shown that Eaf1 serves as an assembly platform for other subunits of NuA4 (14), with structural details missing. A truncation form of Piccolo module has been crystallized and determined at 2.8Å, revealing the inter-subunit contacts of Piccolo module (15). Moreover, a Cryo-EM reconstruction of so-called TEEAA module (Tra1, Eaf1, Eaf5, Actin, Arp4) of NuA4 was resolved at 4.7Å, providing a preliminary insight into the architecture of NuA4 (16).

## Results

### Purification and structure determination of NuA4 complex from *S. cerevisiae*

As one of the two major histone acetyltransferase complexes in *S. cerevisiae*, the NuA4 complex has been extensively studied for almost a couple of decades, however, the structural basis for subunit-subunit interactions, the nucleosome-recognition and acetylation mechanisms remain largely unknown, partially due to the lack of high-resolution structure of NuA4. Here we successfully fused a TAP tag to the C-terminus of Epl1, a subunit of Piccolo module, and subsequently purified endogenous NuA4 complex to homogeneity through tandem-affinity purification (TAP) method (Fig. 1A). The purified NuA4 complex was analysed by SDS-PAGE, and further by mass-spectrometry. Mass-spectrometric analysis confirmed the existence of all subunits of NuA4 complex (Table. S1). To check the catalytic activity of purified NuA4 complex against the nucleosome substrates, NuA4, acetyl coenzyme A (acetyl-CoA), and different forms of nucleosomes were incubated and allowed to react for an hour at 30 _, and immediately subjected to western blot analyse. Our histone acetyltransferase assay showed that NuA4 preferentially acetylases 217-nucleosome (217-NCP), compared to the 147-nucleosome (147-NCP) with shorter extranucleosomal DNA (Fig. 1A), indicating that the catalytic activity of NuA4 is possibly regulated by the chromatin context *in vivo*. To obtain a higher-resolution Cryo-EM structure, stochiometric NuA4 complex was obtained by pooling the peak fractions after performing glycerol gradient centrifugation (Fig. S1A). The sample was concentrated, crosslinked, and then subjected to grid preparation and subsequent Cryo-EM analysis (Fig. S1B and S1C). After multiple rounds of data processing (Fig. S2A), the NuA4 complex was finally resolved at 4 Å (Fig. 1B). Focused classification was further performed for the Core module which doesn’t contain the Lasso-region of Tra1, yielding a reconstruction at 3.8 Å (Fig. 1C, Fig. S2B and S2C). The atomic model of NuA4, built via *de novo* tracing assisted by AlphaFold prediction, fits quite well to the reconstruction maps (Fig. 1D and S3).

**Fig 1.**
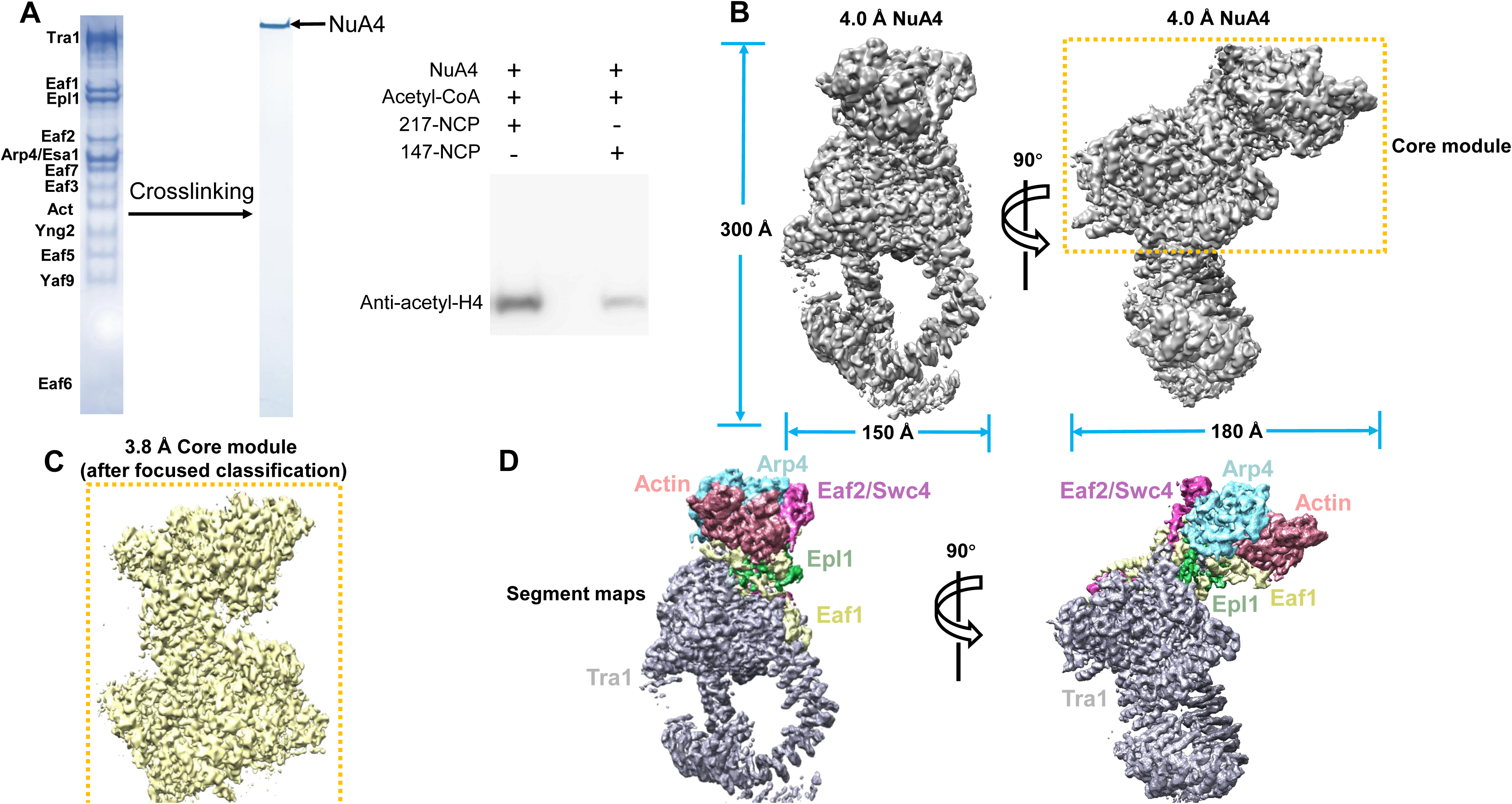
Biochemical and cryo-EM studies of *S. cerevisiae* NuA4 complex. (A) From left to right are the SDS-PAGE bands, crosslinked band of NuA4, and histone acetyltransferase (HAT) assay of NuA4 against different NCP substrates. (B) Electron density map of NuA4 complex and, rotated by 90 °. (C) 3.8 Å electron density map of Core module. (D) Segmentation of the electron density map, rotated by 90 °. Each subunit is indicated and labeled.

### Overall structure of NuA4

Six subunits, including Eaf1, Eaf2/Swc4, Actin, Arp4, Epl1 and Tra1 were apparently seen (Fig. 2A), whereas the TINNTIN module and the other three subunits of Piccolo module, including Esa1, Yng2 and Eaf6 were missing from the map, possibly due to motion. Structural comparison revealed a compositional difference from the published lower-resolution architecture of NuA4, in which Eaf2/Swc4 and Eaf5 were wrongly assigned (16). The Actin-Arp4 and Tra1 modules are connected by the “Neck” region, consisted of the C-terminal domain of Epl1, the post-SANT domain of Eaf2 and the flanking sequences of HSA domain of Eaf1 (Fig. 2A and 2B). The SANT domain of Eaf2, showing a conserved helical feature and whose density was misassigned to Eaf1 in previous study, stacks against the lateral side of Arp4 and therefore stabilizes the Actin-Arp4 module (Fig. 2A).

**Fig 2.**
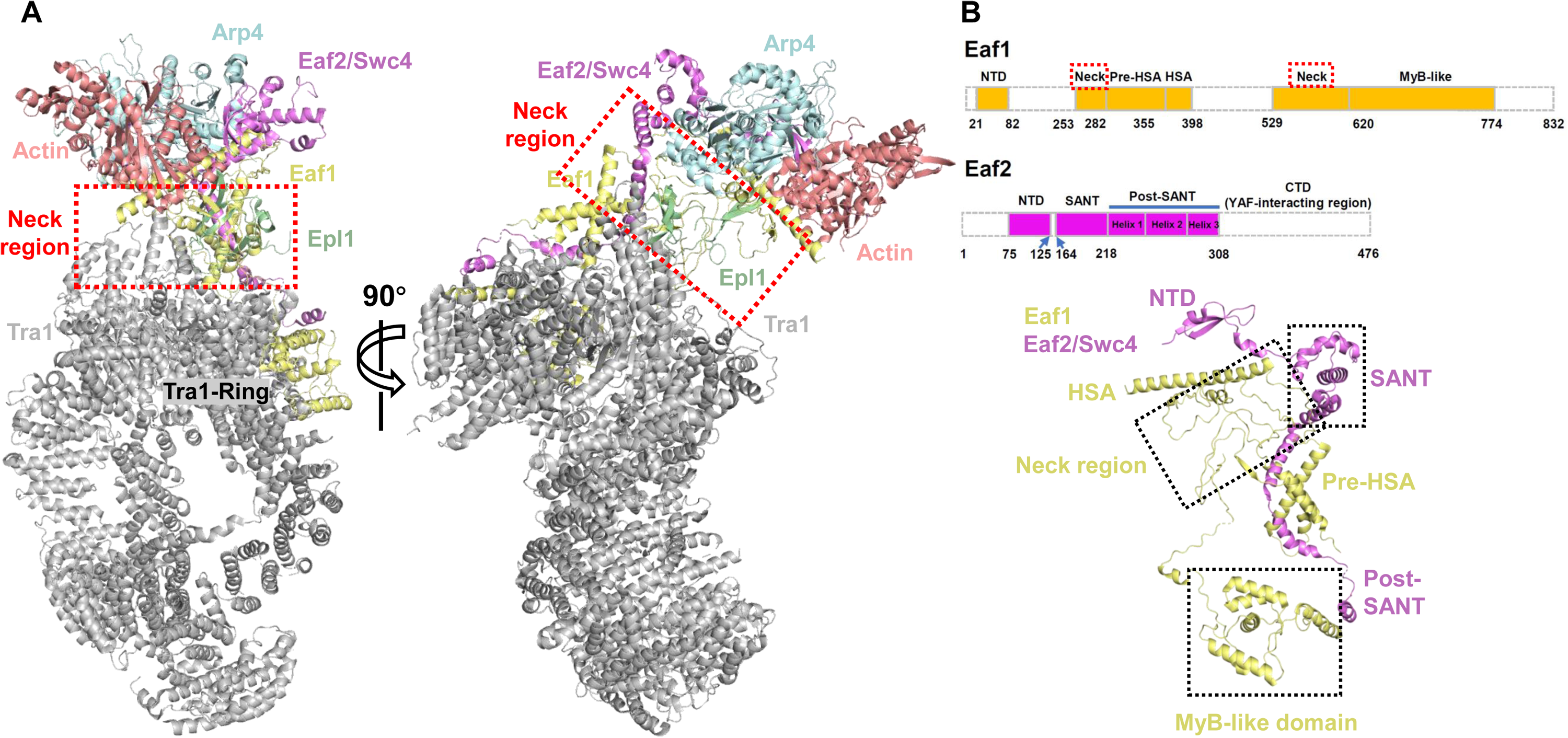
Overall structure of NuA4 complex. (A) Ribbon diagrams of NuA4 subunits, rotated by 90 °. Each subunit is indicated with different color. The “Neck” region is indicated with a red dashed square. (B) From top to bottom are the domain organizations of Eaf1 and Eaf2, and the structure of Eaf1-Eaf2. Residues at domain boundaries are indicated at the bottom.

### Eaf1 and Eaf2 act as the backbone of NuA4 complex

In addition to a previous biochemical study indicating that Eaf1 acts as the backbone of NuA4 complex (14), our structure evidently shows that both Eaf1 and Eaf2/Swc4 adopt long stretched conformations and twist against each other, spanning from the Actin-Arp4 module to the Ring domain of Tra1 module and thereby synergistically forming the backbone of NuA4 complex (Fig. 2A and 2B). According to sequence analysis, Eaf1 comprises a N-terminal domain (Eaf1-NTD), pre-HSA, HSA (Helicase-SANT Associated) and a C-terminal MyB-like domain, whereas Eaf2 consists of a N-terminal domain (Eaf2-NTD), SANT and post-SANT domains (Fig. 2B), all of which are likely essential for their “backbone” roles.

### Actin-Arp4 module is stabilized not only by Eaf1-HSA, but also by Eaf2-NTD

The Actin-Arp4 module is conserved and shared by NuA4, SWR1, INO80 and mammalian PBAF complexes (17-19). The conserved HSA domain resembles a “latch”, stabilizing the Actin-Arp4 module (Fig. 3A). In contrast to the structures of SWR1, INO80 and PBAF complexes, the Actin-Arp4 module in NuA4 is additionally bound by the N-terminal β-sheet of Eaf2, involving several hydrogen-bonds formed by Trp94, Trp78, Asn86, Ser88, Thr90 and Tyr111 from Eaf2, Glu117, Glu125 and Tyr362 from Actin, and Glu135, Trp469 and Glu474 from Arp4 (Fig. 3B). Glu365, Asp373 and several hydrophobic residues of Eaf1-HSA contribute to the interaction between the HSA and the Actin-Arp4 module. Intriguingly, the 644^th^ arginine residue hydrogen bonds with Asp373 of HSA domain, likely stabilizing the HSA position (Fig. 3C, 3D). All the complexes mentioned above have evolved their own HSA domains. Despite the sequence variability, the structures and the binding positions of these HSA domains are, to some extent, evolutionarily conserved (Fig. 3E). Extra density in the nucleotide-binding pocket of Arp4 was observed, and eventually assigned to ATP (Fig. 3F). The ATP binding pocket and the ATP-interacting residues (Fig. 3F) are reminiscent of the structure of Actin-Arp4 module in complex with SWR1-HSA (17). Both structures contain an ATP molecule in the pocket of Arp4, whereas not in that of Actin. We performed a series of HAT assays to confirm whether the ATP binding and hydrolysis would influence the acetylation activity of NuA4. The assays evidently showed that the activity didn’t change either in the presence of ATP, ADP, or ATP-analog AMP-PNP (Fig. 3G), suggesting that the ATP binding and hydrolysis may be not essential for its HAT activity.

**Fig 3.**
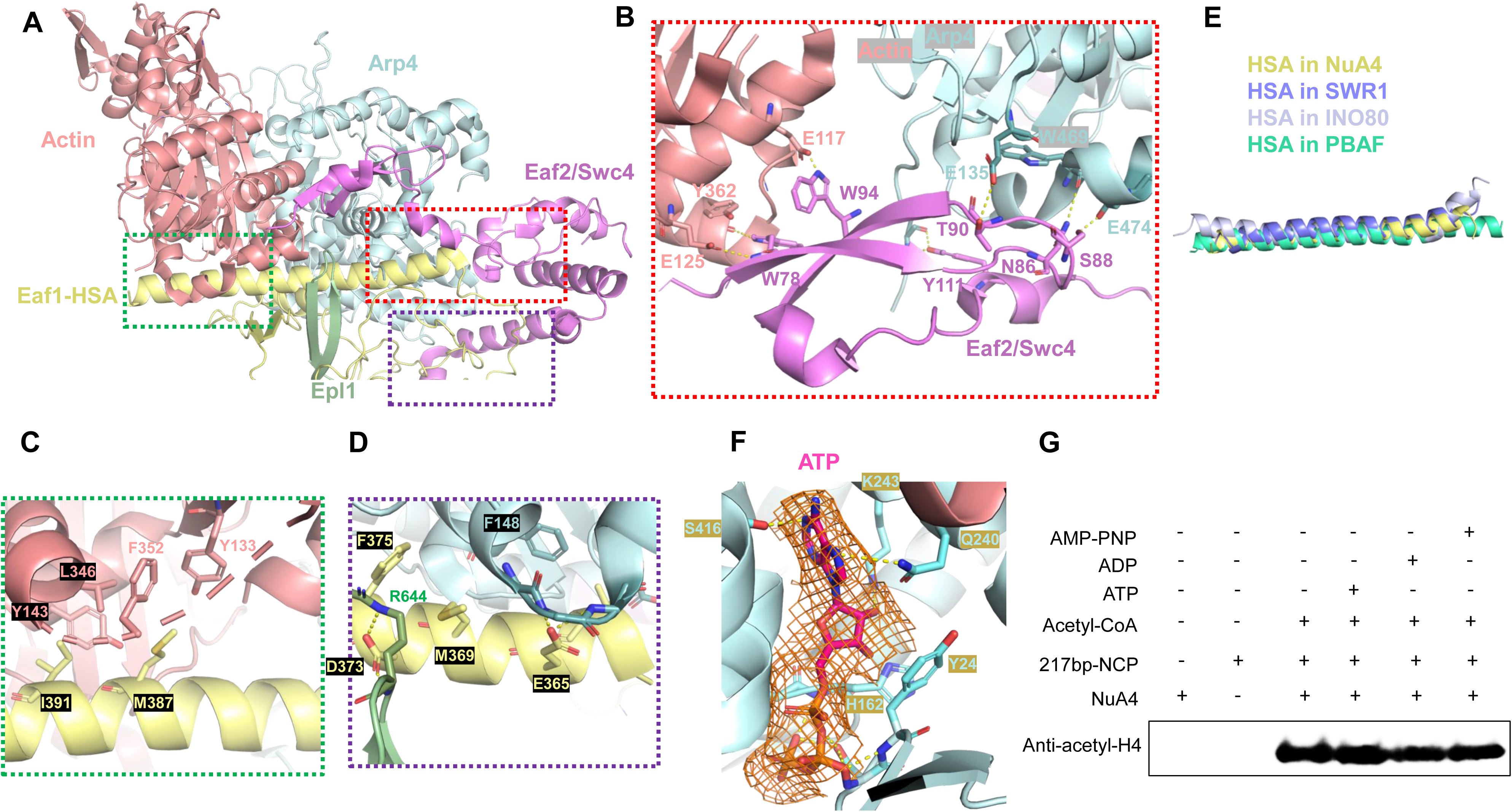
Structures of Actin-Arp4 module in complex with HSA and Eaf2-NTD. (A) Ribbon diagrams of Actin-Arp4-HSA-Eaf2-Epl1. Each subunit is indicated with different color. (B) Structure of Actin-Arp4-Eaf2. The interacting residues are shown as sticks and labeled in different colors. (C) and (D) are the C-terminal and N-terminal interfaces between Eaf1-HSA and the Actin-Arp4 module. The interacting residues are shown as sticks and labeled in different colors. (D) Structure comparison of the HSA domains in NuA4, SWR1, INO80 and PBAF complexes. (E) An ATP molecule and the corresponding electron density are shown as sticks and meshes, respectively. The ATP-interacting residues are shown as sticks and labeled. (F) HAT assay of NuA4 against 217-NCP performed in the presence of acetyl-CoA, in the absence or presence of ATP, ADP and AMP-PNP. Different reaction times are labeled on the bottom of the bands.

### Structure of the “Neck” region and implications for the location of Piccolo module

Previous study reported that the “Neck” region, with most amino acids unassigned, is only formed by Eaf1 subunit (16). After improving the resolution of the “Neck” region in NuA4 to around 3.0-3.2 Å, we came to a different conclusion that the “Neck” region is made up by the post-SANT domain of Eaf2, the C-terminal domain of Epl1 and the HSA-flanking sequences of Eaf1 (Fig. 2A, Fig. 4A). Interestingly, the C-terminal three β-strands of Epl1, whose structures were absent from previous studies, together with the β-strands of Eaf1 constitute two hybrid β-sheets, within which one is sandwich-like. This sandwiched β-sheet, to our knowledge, represents a novel fold of protein complex (Fig. 4B). It is worth noting that the very N-terminal residue of Epl1 in the map is Thr539. Given that the N-terminal region of Epl1 (Thr539-preceding sequence) interacting with Esa1 and therefore participating in Piccolo module formation (15), it is possible to indicate the location of the remaining part of Piccolo module (Fig. 4B). This indicated location of Piccolo module may also shed light on the nucleosome binding position on NuA4.

**Fig 4.**
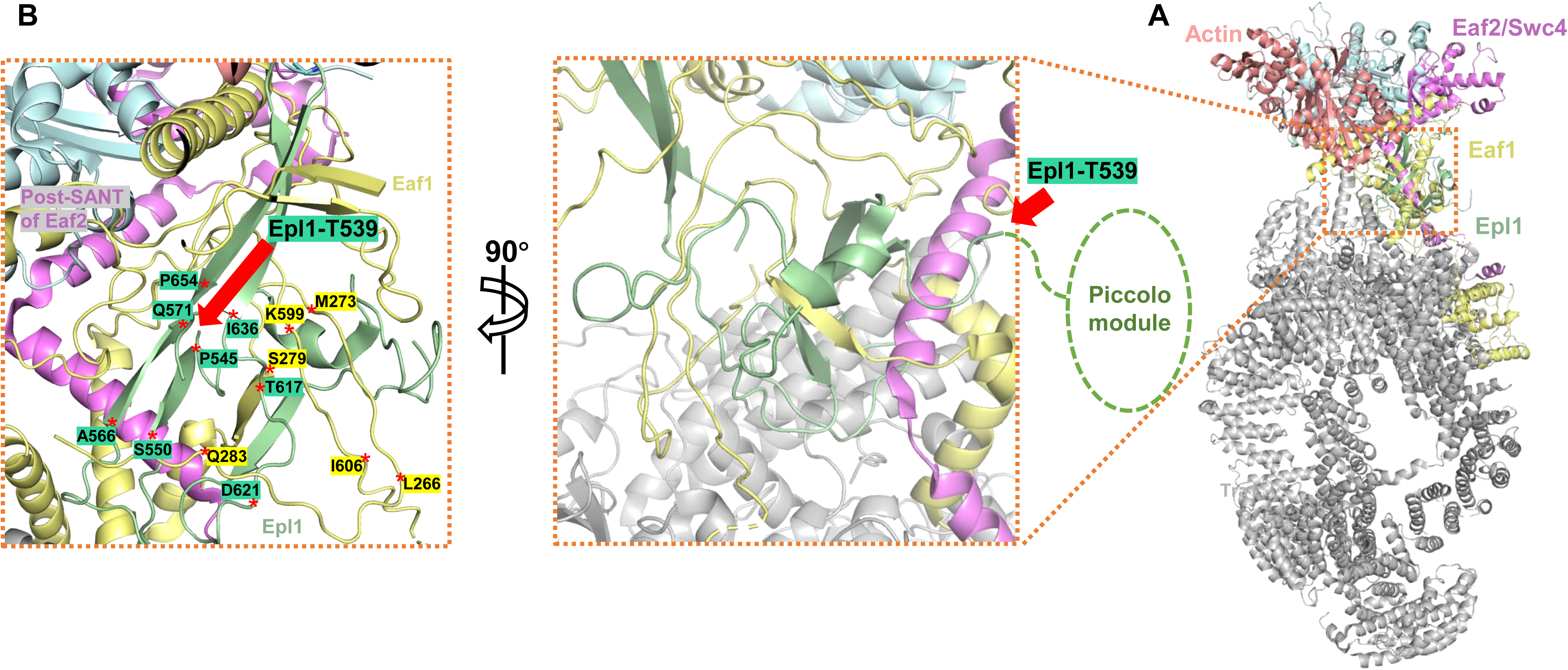
Structures of the “Neck” region. (A) Ribbon diagrams of NuA4. The “Neck” region is indicated with a salmon dashed square. (B) Expanded view of the “Neck” region formed by Eaf1, Eaf2 and Epl1, rotated by 90 °. Residues at the β-strand boundaries are indicated and labeled with different backgrounds. The amino acid T539 of Epl1 is indicated with a red arrow, and the possible position of Piccolo module is indicated with a green circle.

### An unexpected position of Yaf9 in NuA4 complex

As described above, the Eaf2 subunit spans from the Arp4 protein to the Ring domain of Tra1, with its three α-helices of post-SANT domain tethering Eaf1 and Tra1 proteins. These α-helices were designated “tethering helix 1”, “tethering helix 2” and “tethering helix 3”, respectively. The C-terminal region following the tethering helix 3 was missing from the reconstruction map. Considering the C-terminus of Eaf2 is required for interaction with Yaf9 (14), a histone-tail binding protein of NuA4, and the binary structure of Eaf2-Yaf9 could be predicted by Alphafold-Multimer, we set out to unveil the location of Yaf9 on NuA4. Despite interference and a slight bending of the tethering helix 1 and 2, comparing to a long-merged tethering helix in predicted structure, the overall conformation and the tethering helix 3 are similar to those of predicted one (Fig. 5A, 5B). The predicted Eaf2-Yaf9 structure demonstrated that the C-terminal domain (324 aa to 476 aa) forms a stable complex with Yaf9 (Fig. 5B). Taken the length of the linker between the tethering helix 3 and the C-terminal domain of Eaf2 into consideration, the Yaf9 location could be roughly indicated to be adjacent to the Ring domain of Tra1, instead of the previously proposed position around Actin-Arp4 module.

**Fig 5.**
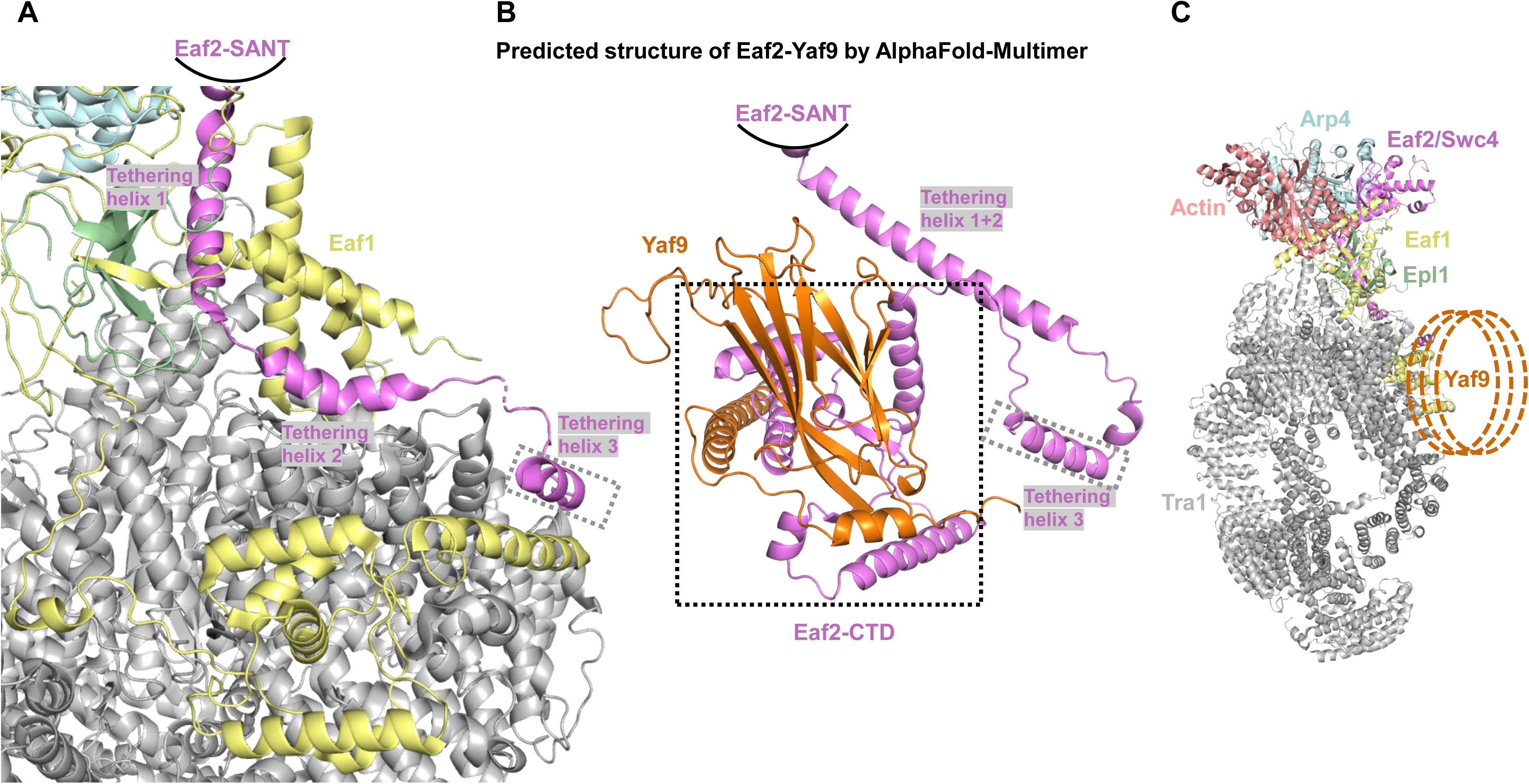
A possible binding position of Yaf9. (A) Ribbon diagrams of the interaction among the post-SANT domain of Eaf2, Eaf1 and Tra1. Each subunit is indicated with different color, and the tethering helices are labeled. (B) The predicted structure of Eaf2-Yaf9 by AlphaFold-Multimer. The tethering helices are labeled and the C-terminal domain of Eaf2 is indicated with a black dashed square. (C) The position of Yaf9 in NuA4. The possible positions of Yaf9 are indicated with orange dashed circles.

### A proposed model of nucleosome-recognition and acetylation mechanism of NuA4

Component missing in the Cryo-EM map prompted us to reconstitute a NuA4-nucleosome complex for further studies. The separately purified NuA4 and 217-nucleosome were mixed at a molar ratio of 1:1 and subjected to glycerol gradient sedimentation. NuA4 complex didn’t bind 217-NCP *in vitro*, unexpectedly, as two proteins migrated at clearly different positions (Fig. 6A), though NuA4 exhibited high activity against the nucleosome substrate (Fig. 3G). We then overexpressed the proteins of Piccolo module through baculovirus system, and further purified to homogeneity (Fig. 6B). Gel-shift and HAT assays showed that the Piccolo module alone is capable of binding and acetylating nucleosome substrate, comparing to the entire NuA4 complex (Fig. 6B and Fig. 6C). Combining the structural and biochemical data together gave rise to a nucleosome-recognition and acetylation model for NuA4 complex. In this model, the Piccolo module alone is able to bind and acetylate histone tail of nucleosome, while entire NuA4 complex adopts an auto-inhibited conformation, likely burying its Piccolo module into other regulatory subunits. Through an unknown mechanism, the Piccolo module becomes accessible, and the entire complex turns to be activated upon additions of nucleosome and acetyl-CoA. This process possibly involves structural rearrangement of NuA4 induced by the substrate and the co-factor acetyl-CoA, which in turn restores the catalytic activity of NuA4 and leads to hyper-acetylation (Fig. 6D).

**Fig 6.**
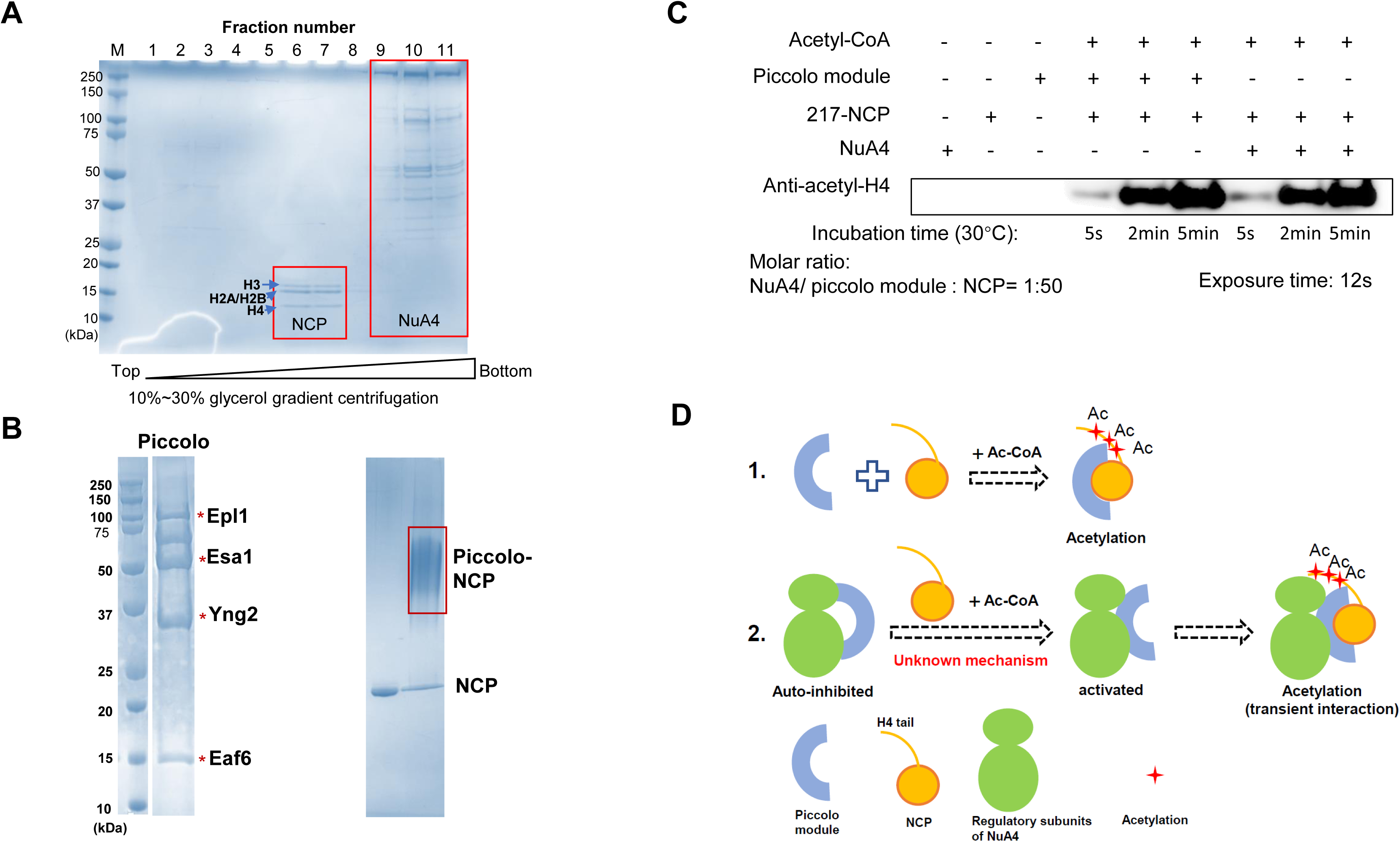
A proposed model for nucleosome recognition and acetylation of NuA4. (A) Glycerol gradient sedimentation of 217-NCP-NuA4. Gradients were fractionated in 500 μl, loaded on a 12% SDS-PAGE gel and visualized by Coomassie blue staining. The peak fractions of NuA4 and NCP migrated at different positions are indicated with red squares, respectively. The histone proteins are indicated with blue arrows and labeled. (B) From top to bottom are the SDS-PAGE analysis of purified Piccolo module and the gel-shift result of Piccolo module against nucleosome, visualized by Coomassie blue staining. (C) Comparison of the HAT activity of NuA4 and its Piccolo module. The HAT reactions were quenched at time points of 5 seconds, 2 and 5 minutes, respectively. The exposure time for NuA4 and Piccolo module 12 seconds. (D) A proposed model to illustrate the nucleosome recognition and acetylation mechanisms of NuA4.

## Discussion

In eukaryotes, both nucleosome-modifying enzymes and remodelers are critical for transcription regulation. Thus far, high-resolution structures of most remodelers complexed with nucleosome have been reported. The stable complexes of remodeler-nucleosome could be captured by introducing an ATP analog ADP-BeF6 into the system and then be used for structural studies. Different from those remodelers, NuA4 doesn’t need ATP or its hydrolysis to perform its catalytic activity (Fig. 3G). It is worth mentioning that NuA4 exhibits strikingly strong activity, as a complete reaction could be achieved within two minutes (Fig. 6C), making it difficult to reconstitute the NuA4-nucleosome complex *in vitro*. According to our proposed model, the NuA4 upon structural rearrangements transiently interacts with and rapidly acetylases its substrate. The auto-inhibited conformation that adopted by apo-NuA4, and nucleosome and/or acetyl-CoA induced structural rearrangements may be significant for substrate specificity of NuA4. Beyond Cryo-EM, other tools such as single molecule method may be more useful to investigate the dynamics and kinetics of the NuA4-mediated acetylation process.

Intriguingly, NuA4 and SAGA, two major histone acetyltransferase complexes in *S. cerevisiae*, share the largest subunit Tra1 (11). The structure of Tra1 in NuA4 was superimposed to that in SAGA. We found the Tra1 structures showed considerable similarity, with a r.m.s.d (root mean square deviation) of 1.002 Å over 3404 Cα atoms. The NuA4-specific subunits rotates around 90 ° with respect to the SAGA-specific proteins (Fig. S4). It is very interesting that there is little steric clash between the NuA4-specific and SAGA-specific subunits. It remains interesting that why the *S. cerevisiae* and other fungal organisms didn’t evolve a super complex, within which the NuA4-specific subunits assembled on one side of Tra1 and the SAGA-specific subunits assembled on the perpendicular side. After careful check, we obtained the answer. We found the C-terminal MyB-like domain of Eaf1 binds to the assembly surface of SAGA-specific subunits (Fig. S4), which may preclude the sequential assembly of these subunits on Tra1 of NuA4, maintaining the uniqueness of NuA4.

### Data availability

Cryo-EM maps for Core module of NuA4, and entire NuA4 complexes have been deposited in the Electron Microscopy Data Bank (EMDB) under accession codes EMD-33794 and EMD-33796. Coordinates for the models have been deposited in the Protein Data Bank (PDB) under accession numbers 7YFN and 7YFP, respectively.

## Supporting information

Supplemental materias

## Acknowledgements

We are grateful to Ms. Chengqian Zhang for mass spectrometric analysis. We thank the Bio-Electron Microscopy Facility of ShanghaiTech University for assistance in data collection. The research was supported by a grant from ShanghaiTech University.

## Author contributions

H.Z., and R.D.K. designed the experiments; L.J., and L.Z. carried out the experiments.; K.X., and H.Z. collected and processed the cryo-EM data.; H.Z. built the models.; H.Z. and R.D.K. wrote the paper.

