## Supplemental materias for "Structure of the histone acetyltransferase NuA4 complex"

### Materials and methods

#### Purification of yeast NuA4 complex and its Piccolo module

40 liters of *S. cerevisiae* cells with Epl1 bearing a C-terminal TAP tag were grown at 30 °C and harvested at $OD_{600} \approx 10$ by centrifugation. Cell pellets were resuspended in lysis buffer A (100 mM HEPES, pH 8.0, 600 mM NaCl, 6 mM DTT and 2X EDTA-free protease inhibitor cocktail), and then homogenized to prepare the whole-cell extraction. To get rid of contaminated nucleic acid, 0.25% (v/v) PEI precipitation was applied. The supernatant containing the protein of interest was further precipitated using 50% (v/v) ammonium sulfate, and the precipitation was dissolved with buffer B (50 mM HEPES, pH 8.0, 25 mM ammonium sulfate, 3 mM DTT and 1X EDTA-free protease inhibitor cocktail). The solute was further cleared by centrifugation (8,000 rpm, 4 °C, 90 minutes) and the supernatant was then pumped through IgG Sepharose-6 Fast Flow resin (GE Healthcare) at 4 °C overnight. After extensively washing with buffer containing 50 mM HEPES, pH 8.0, 150 mM ammonium sulfate and 3 mM DTT, overnight on-column cleavage was performed by adding TEV protease into the IgG resin. The eluate was then concentrated and subjected to 10% to 30% glycerol gradient centrifugation (40,000 rpm, 4 °C, 10 hours). The peak fractions were pooled, dialyzed against buffer C (25 mM HEPES, pH 8.0, 200 mM NaCl and 3 mM DTT) and concentrated to approximately 1 mg/ml before grid preparation. To protect NuA4 complex from falling apart, the concentrated sample was crosslinked by 0.1% (v/v) glutaraldehyde in the presence of 0.01% NP40 (v/v) on ice for 10 minutes before grid freezing.

The genes encoding the subunits of Piccolo module were amplified from the genomic DNA of *S. cerevisiae* and cloned into pFastBac-1 vector with a 6× His tag at the N-terminus of Epl1 and a Flag tag at the C-terminus of Esa1. Each baculovirus was produced using the Bac-to-Bac baculovirus expression system (Invitrogen). The Piccolo module was overexpressed in High Five cells by co-infection of four baculovirus for 48 hours at 27 °C. One liter of High Five cells ($2.0 \times 10^6$ cells $ml^{-1}$ cultured in ESF921 medium) was harvested, centrifuged. The Cell pellets were resuspended with buffer containing 25 mM HEPES, pH 8.0, 300 mM NaCl and 3mM DTT, and lysed by sonication. The lysate was cleared by centrifugation, purified using Ni-NTA (Qiagen) and anti-Flag columns (Genscript), and further cleaned using Heparin and size-exclusion chromatography (Superdex 200, GE Healthcare) in buffer containing 25 mM HEPES, pH 8.0, 100 mM NaCl and 3 mM DTT. The peak fractions were concentrated to ~1mg/ml before use.

Vitrobot Mark IV was used for grid plunging. 300-mesh Quantifoil R1.2/1.3 grids were glow-discharged before grid freezing. 3 μl of crosslinked NuA4 complex was applied to the grid. After incubation for 60 seconds, the grid was blotted for 2.5 seconds before being plunged into liquid ethane with 100% chamber humidity at 8 °C. All grids were stored in liquid nitrogen until data collection.

The *Xenopus* histone octamer was overexpressed, purified, and reconstituted with 147 bp and 217 bp Widom-601 DNAs as described previously (1).

#### Cryo-EM data collection and image processing

Automated data acquisitions were performed using SerialEM (2) on a Titan Krios equipped with a Gatan K3 Summit direct electron detector (Gatan, Inc.) and operating at 300 kV with a nominal magnification of 18,000 ×. All images were automatically recorded in the super-resolution mode, with a defocus range from 1.3 to 2.0 μm. Each movie stack was dose-fractionated to 40 frames with a total electron dose of ~60 $e^-/Å^2$ and a total exposure time of ~3.4 seconds. Movie stacks were motion-corrected and the defocus value was estimated using the modules in cryoSPARC (3). The particles were picked, extracted, and then subjected to cryoSPARC (3) for 2D classification, heterogeneous and non-uniform (NU) refinements, yielding a reconstruction of NuA4 at 4 Å. To improve the resolution of the core module, a mask around the core module was generated and used for particle subtraction in cryoSPARC (3). Heterogeneous refinement and local refinement were applied, resulting to a 3.8 Å reconstruction of the Core module.

**Model building**

All the model buildings were performed using the cryo-EM module of Phenix package (4), Chimera (5), and COOT (6).

The 3.7 Å atomic model of Tra1 (PDB entry: 5OJS) (7) was docked into our 3.8 Å Cryo-EM map using Chimera (5), and then manually adjusted in COOT (6).

The long and flexible linkers in AlphaFold-predicted structures of Actin and Arp4 were first delated using COOT (6), and then docked into the 3.8 Å Cryo-EM map of Core module using Chimera (5).

The remaining parts of the cryo-EM maps were successfully assigned to three subunits including Epl1, Eaf1 and Eaf2, owing to the relatively higher resolution of these regions and the consistencies between the map features and the AlphaFold-predicted structures.

All above-mentioned separate models were merged, and then subjected to Phenix for several rounds of real-space refinement. All the structure figures were generated by PyMOL (https://pymol.org/2/) and Chimera (5).

**Histone acetyltransferase assay**

All the reactions mentioned in the manuscript were performed in reaction buffer containing 25 mM HEPES (pH 8.0), 50 mM ammonium sulfate, 3 mM DTT, 100 μM acetyl-CoA, and 1 μM NCP (217-NCP or 147-NCP). To test the effects of different nucleotides on the reaction, 0.1 μM NuA4 and equal molar of the corresponding nucleotides were added to the reaction buffer and incubated at 30 °C, respectively. To compare the catalytic activity of NuA4 and Piccolo module, 20 nM NuA4 and an equal molar of Piccolo proteins were added to the reaction buffer, respectively, and then were allowed to react for 5 seconds, 2 minutes and 5 minutes at 30 °C, respectively. All the reactions were quenched by adding 5X SDS loading buffer and heated to 100 °C for 5 minutes. The reaction mixtures were loaded to a 12% SDS-PAGE gel, and subsequently analyzed using the primary antibody against acetylated-lysine of H4 (Abcam). The signals were quantified by ChemiDoc (Bio-Rad).

**Gel-shift assay**

To test the interaction, Piccolo module of NuA4 was mixed with an equal molar of NCP at a for an hour on ice, and then were loaded on a 3-12% Native-PAGE gel and detected by Coomassie blue.

**Glycerol gradient centrifugation**

Purified NuA4 and an equal molar of 217-NCP were incubated for 1 hour on ice in the buffer containing 25 mM HEPES (pH 8.0), 50 mM ammonium sulfate, 3 mM DTT and 5 mM $MgCl_2$. The mixture was applied to a 10–30% glycerol gradient in the same buffer and ultracentrifuged at 40,000 rpm (SW41 rotor) for 7 hours at 4 °C. Gradients were fractionated in 500 μl and analysed with SDS-PAGE. The gels were stained with Coomassie blue.

**Mass-spectrometry**

The NuA4 sample was released from SDS-PAGE gel, and then were reduced, alkylated and digested by trypsin (Promega, V5111, USA). The obtained peptides were desalted with C18 ZipTip (Millipore, USA), and then vacuum dried. Peptides were dissolved in 0.1% FA for separating and analyzing by label-free quantification on an Easy-nLC 1000 system coupled to a Q Exactive HF (Thermo Scientific, USA). MaxQuant (v.1.6.5.0) was used to process the raw data of LC-MS/MS analysis according to the *S. cerevisiae* Uniprot FASTA database (release 2020_12, 6, 050 sequences). In the parameters setting, analysis allowed two missed cleavage sites of trypsin. Mass error of precursor ions was set to 5 ppm, and fragment ions was set to 0.02 Da. The fixed modification selected carbamidomethylation on Cys and variable modifications set as oxidation (M), deamidation (NQ), acetylation (Protein N-term). False discovery rate (FDR) thresholds were set at 1%. Minimum peptide length allowed by the analysis was 7.


1. Xu TH*, et al.* (2020) Structure of nucleosome-bound DNA methyltransferases DNMT3A and DNMT3B. *Nature* 586(7827):151-+.
2. Mastronarde DN (2005) Automated electron microscope tomography using robust prediction of specimen movements. *J Struct Biol* 152(1):36-51.
3. Punjani A, Rubinstein JL, Fleet DJ, & Brubaker MA (2017) cryoSPARC: algorithms for rapid unsupervised cryo-EM structure determination. *Nat Methods* 14(3):290-+.
4. Adams PD*, et al.* (2002) PHENIX: building new software for automated crystallographic structure determination. *Acta Crystallogr D* 58:1948-1954.
5. Pettersen EF*, et al.* (2004) UCSF chimera - A visualization system for exploratory research and analysis. *J Comput Chem* 25(13):1605-1612.
6. Emsley P & Cowtan K (2004) Coot: model-building tools for molecular graphics. *Acta Crystallogr D* 60:2126-2132.
7. Diaz-Santin LM, Lukoyanova N, Aciyan E, & Cheung ACM (2017) Cryo-EM structure of the SAGA and NuA4 coactivator subunit Tra1 at 3.7 angstrom resolution. *Elife* 6.


**Figure legends**

**Fig S1. Purification and Cryo-EM analysis of NuA4.**

(A) SDS-PAGE analysis of NuA4 fractions after glycerol gradient centrifugation. The fractions were loaded on 4-12% resolving gels and visualized by Coomassie blue staining. The peak fractions chosen for cryo-EM study are indicated with a red dashed square.

(B) A representative electron micrograph of NuA4 complex. Scale bar, 500 Å.

(C) A representative 2D class averages of NuA4 particles.

**Fig S2. Cryo-EM analysis of NuA4 complex.**

(A) A flow-chart for cryo-EM data processing of NuA4 complex.

(B) Quality of the cryo-EM maps. From top to bottom are the viewing direction distribution plot, the gold standard FSC curves, and the local resolution map for each map, respectively.

**Fig S3. Fitting the built models of Eaf1, Eaf2, Epl1, Tra1 and the Actin-Arp4 module to the EM density.**

Fittings of Eaf1, Eaf2, Epl1 and Actin-Arp4 to their density maps were done in PyMOL, and the fitting of Tra1 to its density map is achieved using Chimera.

**Fig S4. Structure comparison of NuA4 and SAGA complexes.** SAGA (PDB entry: 6T9I) is colored in gray and NuA4 is colored in yellow. The right panel is an expanded view of the interface between Tra1 and SAGA-specific subunits. The MyB-like domain of Eaf1 is red-colored and labeled.

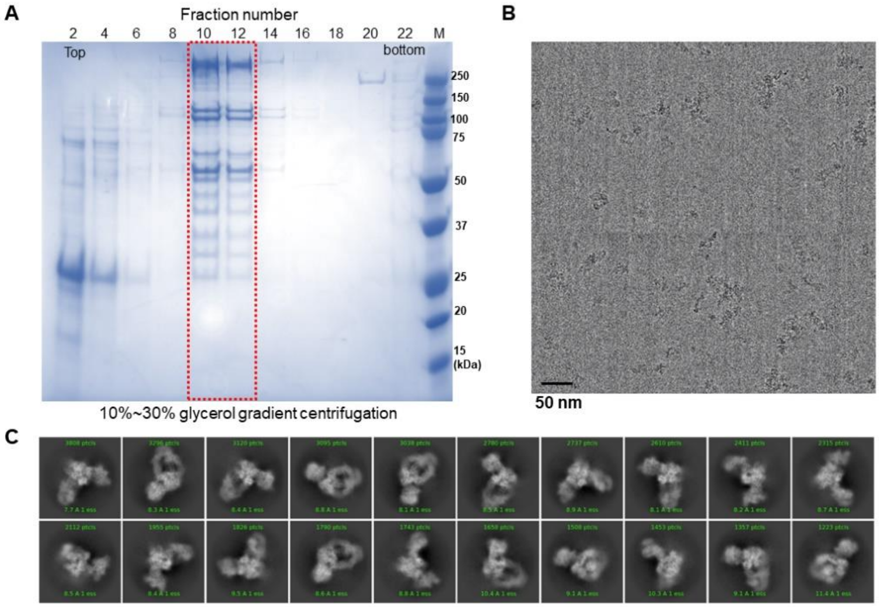


Fig. S1

Fig. S2

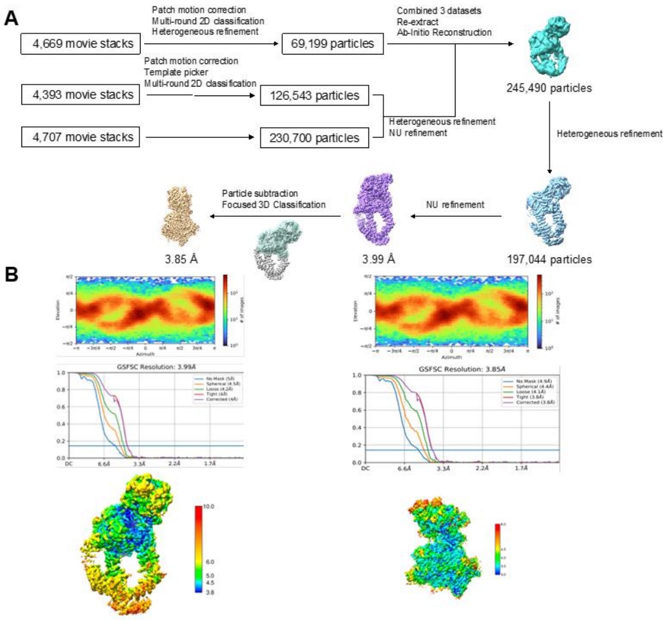
A
4,669 movie stacks
Patch motion correction
Multi-round 2D classification
Heterogeneous refinement
69,199 particles
Combined 3 datasets
Re-extract
Ab-Initio Reconstruction
245,490 particles
4,393 movie stacks
Patch motion correction
Template picker
Multi-round 2D classification
126,543 particles
4,707 movie stacks
230,700 particles
Heterogeneous refinement
NU refinement
Heterogeneous refinement
197,044 particles
NU refinement
3.99 Å
Particle subtraction
Focused 3D Classification
3.85 Å
B
GSFSC Resolution: 3.99Å
GSFSC Resolution: 3.85Å

Fig. S3

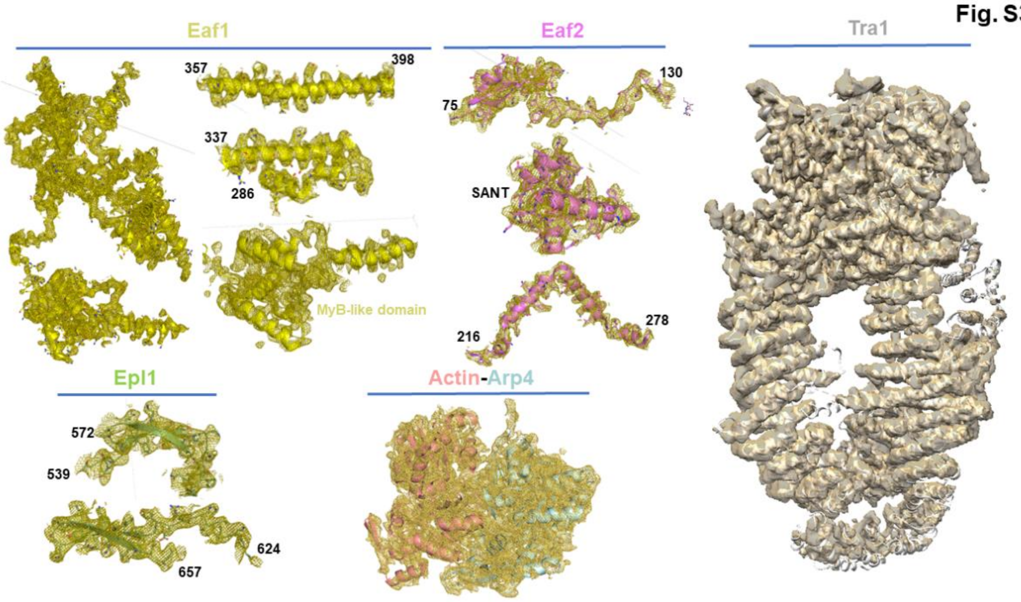
Eaf1
357
398
337
286
MyB-like domain
Eaf2
130
75
SANT
216
278
Tra1
Epl1
572
539
624
657
Actin-Arp4

Fig. S4

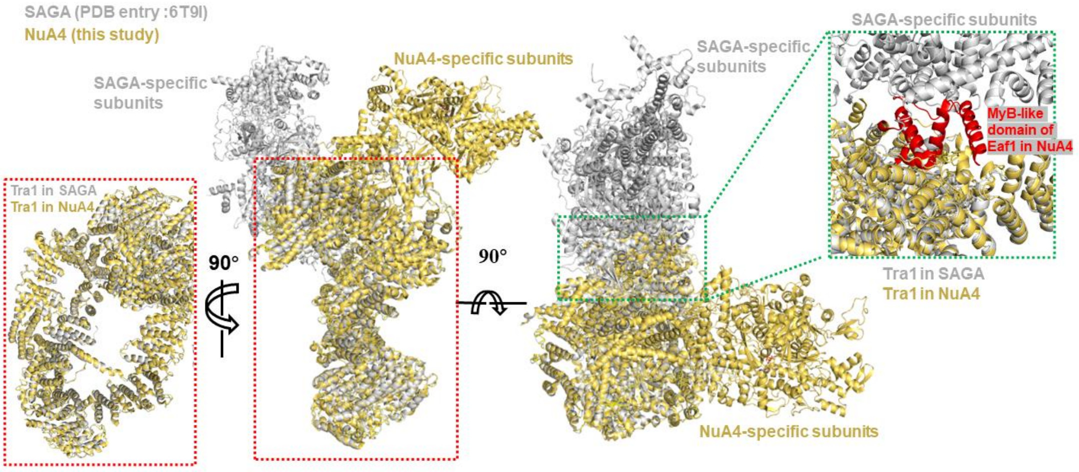


### Table S1. Mass-spectrometric analysis of the purified NuA4 complex from *S. cerevisiae*

| Gene names | Unique peptides | Sequence coverage (%) | Score | Intensity | Mol. weight (kDa) |
|---|---|---|---|---|---|
| TRA1 | 232 | 53 | 323.31 | 3.8084E+11 | 433.17 |
| EAF1 | 99 | 72.3 | 323.31 | 1.0024E+11 | 112.5 |
| EPL1 | 74 | 70 | 323.31 | 48199000000 | 96.737 |
| SWC4/EAF2 | 46 | 62 | 323.31 | 26937000000 | 55.212 |
| ESA1 | 41 | 63.8 | 323.31 | 31295000000 | 52.612 |
| EAF3 | 32 | 73.8 | 166.47 | 16343000000 | 45.203 |
| ARP4 | 31 | 42.7 | 274.06 | 16187000000 | 54.831 |
| EAF7 | 30 | 60.2 | 165.66 | 9514700000 | 49.391 |
| ACT | 25 | 69.6 | 175.19 | 19942000000 | 41.689 |
| YNG2 | 21 | 55.3 | 136.27 | 9069400000 | 32.086 |
| EAF5 | 19 | 60.9 | 323.31 | 2271300000 | 31.644 |
| YAF9 | 19 | 73.5 | 104.35 | 9453200000 | 25.981 |
| EAF6 | 9 | 85.8 | 4.9285 | 353420000 | 12.895 |

**Table S2. Cryo-EM data collection, refinement, and validation statistics.**

| Complex/subcomplex | Core module | Entire NuA4 |
|---|---|---|
| EMDB | EMD-33794 | EMD-33796 |
| PDB | 7YFN | 7YFP |
| **Data collection and processing** | | |
| Magnification | 18,000 | 18,000 |
| Voltage (kV) | 300 | 300 |
| Electron exposure ($e^-/Å^2$) | 60 | 60 |
| Defocus range (-µm) | 1.3-2.0 | 1.3-2.0 |
| Pixel size (Å) | 0.66 (super-resolution mode) | 0.66 (super-resolution mode) |
| Symmetry imposed | C1 | C1 |
| Initial particle images (no.) | 426,442 | 426,442 |
| Final particle images (no.) | 197,044 | 197,044 |
| Map resolution (Å) | 3.8 | 4.0 |
| FSC threshold | 0.143 | 0.143 |
| Map resolution range (Å) | 3.0-5.0 | 3.1-5.2 |
| **Refinement** | | |
| Initial model used | *de novo* modelling assisted by AlphaFold2 | 5OJS |
| Model resolution (Å) | 3.8 | 4.0 |
| FSC threshold | 0.143 | 0.143 |
| Model resolution range (Å) | 3.6-3.8 | 3.8-4.1 |
| Map sharpening B factor ($Å^2$) | | |
| Model composition | | |
| Non-hydrogen atoms | 25,868 | 40,390 |
| Protein residues | 3,222 | 5,009 |
| Ligands | 1 | 1 |
| B factor ($Å^2$) | | |
| Protein | 111.2 | 203.5 |
| Ligand | 5.6 | 53.4 |
| R.m.s. deviations | | |
| Bond lengths (Å) | 0.007 | 0.008 |
| Bond angles (°) | 1.14 | 1.16 |
| Validation | | |
| MolProbity score | 2.84 | 2.62 |
| Clashscore | 14 | 22 |
| Poor rotamers (%) | 0.3 | 0.4 |
| Ramachandran plot | | |
| Favored (%) | 90.7 | 90.2 |
| Allowed (%) | 8.9 | 9.3 |
| Disallowed (%) | 0.4 | 0.5 |
